# AMaLa: Analysis of Directed Evolution Experiments via Annealed Mutational approximated Landscape

**DOI:** 10.1101/2021.07.26.453757

**Authors:** Luca Sesta, Guido Uguzzoni, Jorge Fernandez-de-Cossio Diaz, Andrea Pagnani

## Abstract

We present Annealed Mutational approximated landscape (AMaLa), a new method to infer fitness landscapes from Directed Evolution experiment sequencing data. Directed Evolution experiments typically start from a single wild-type sequence, which undergoes Darwinian in vitro evolution acted via multiple rounds of mutation and selection with respect to a target phenotype. In the last years, Directed Evolution is emerging as a powerful instrument to probe fitness landscapes under controlled experimental condition and, thanks to the use of high-throughput sequencing of the different rounds, as a relevant testing ground to develop accurate statistical models and inference algorithms.

Fitness landscape modeling strategies, either use as input data the enrichment of variants abundances and hence require observing the same variants at different rounds, or they simply assume that the variants at the last sequenced round are the results of a sampling process at equilibrium. AMaLa aims at leveraging effectively the information encoded in the time evolution of all sequenced rounds. To do so, on the one hand we assume statistical sampling independence between sequenced rounds, and on the other we gauge all possible trajectories in sequence space with a time-dependent statistical weight consisting of two contributions: (i) a statistical energy term accounting for the selection process, (ii) a simple generalized Jukes-Cantor model to describe the purely mutational step.

This simple scheme allows us to accurately describe the Directed Evolution dynamics in a concrete experimental setup and to infer a fitness landscape that reproduces correctly the measures of the phenotype under selection (e.g. antibiotic drug resistance), notably outperforming widely used inference strategies. We assess the reliability of AMaLa by showing how the inferred statistical model could be used to predict relevant structural properties of the wild-type sequence, and to reproduce the mutational effects of large scale functional screening not used to train the model.

## I. INTRODUCTION

Over the last years, the development of increasingly accurate high-throughput biochemical assays jointly with massive parallel sequencing techniques has established large-scale genetic screening as a fundamental tool for the investigation of the relationship between evolution, fitness and other important biological concepts that were behind experimental research. [1–27].

These experiments that simultaneously screen up to millions of variants of a given protein come into two main flavors. Deep Mutational Scanning (DMS) experiments [8], where all the combinatorial complexity of the experiment is encoded in the initial library that undergoes several iterative steps of target selection and amplification. In contrast, in Directed Evolution experiments [28], at each round of the procedure, mutations are randomly created with a tuned mutation rate by error-prone PCR (ep-PCR). In either case, the typical pipeline sees a library of protein variants undergoing cycles of selection for functional activity. At some of the intermediate steps, a sample of the mutants population is sequenced to assess the relative variant abundances.

Recently, high-throughput screening experiments have been used to predict the folded three-dimensional structure [26, 29]. It has been argued that screening experiments without mutations, like DMS, are not able to probe the sequence space deeply enough to generate a statistically relevant signal of the folded structure. In both studies [26, 29], the author’s experimental solution involves the Directed Evolution framework introducing an error-prone PCR to explore a broader sequence space region. Then, their idea is to apply Direct Coupling Analysis (DCA) to the artificially generated protein variants by the Directed Evolution experiment. DCA was originally developed to model co-evolution in homologous protein families [30, 31]. It uses the inferred epistatic interaction between pairs of residues to reconstruct the contact map. In this strategy, a maximum entropy model is learned from the outcome of the last round of selection. This set of proteins is the output of a functional selection constrained to the structural properties of the wild-type, mimicking the protein families generated by natural evolution. Some of the differences between the two processes involve the time duration, the broader sequence diversity, the population sizes, the mutation rates, the variability in the cellular environments where the protein operates (e.g different temperatures), among others.

Besides the motivation to predict the protein structure, having a correct model of the genotype-phenotype association permits answering fundamental questions about the relationship between fitness landscape and molecular evolution ([32–34]) and on a more practical side to design novel effective proteins ([35, 36]).

As more high-throughput sequencing data of screened libraries are available, new computational methods for accurate statistical modeling of the genotype-phenotype association are actively developed. Most of the computational strategies developed so far rely on two approaches: (i) DCA-inspired models of phylogenetically related sequences used to describe local fitness landscapes [37–44], and very recently in [45]. (ii) supervised machine learning approach on sequenced sample of high-throughput functional assays or screening experiments. In this case, a statistical model of the mutants’ fitness is inferred from a subset of the sequencing data (training set) with machine learning techniques developed to solve a specific – generally non-linear – regression problem [26, 27, 41, 46–49].

More recently, alternative unsupervised strategies have been proposed [50, 51] to cope with all sequencing information coming from screening experiments. In [51] a probabilistic model is described, that takes into account three different steps always occurring in screening experiments: i) selection, ii) amplification, and iii) sequencing. Although such models are very effective to describe the Deep Mutational Scanning experiments in absence of the mutagenesis step, it relies on the variations of the variants’ relative abundances across rounds and hence of the sample of the same variants at different time-steps with sufficient statistics. However, several screening setups and in particular Directed Evolution experiments do not allow for this computation. It can be due to a small sequencing depth (related to the initial library size) or to an additional error-prone PCR step to create more variability as in the Directed Evolution experiment. In the last case, new sequences appear at each round of the experimental pipeline, thus the variation of the library composition is not due only to fitness selection but also to a stochastic drift.

In this paper, we propose Annealed Mutational approximated Landscape analysis (AMaLa), a new unsupervised inference framework that effectively takes into account the mutational aspects of Directed Evolution experiments in terms of a global likelihood to observe a time series of variants’ abundances. AMaLa specifically aims to model the whole in-vitro evolutionary trajectory, contrarily to the DCA approach that considers only the last sequenced round. The genotype-to-phenotype mapping is defined in terms of multivariate Potts-like hamiltonians with both additive contribution from individual residues and pairwise epistatic contributions. We remark that the method does not require computing an enrichment ratio of the variants population.

The effectiveness of AMaLa is assessed by analyzing three Directed Evolution experiments [26, 29] in terms of the ability: (i) to predict the mutation effect on the fitness assessed via large-scale functional screening, (ii) to reconstruct the contact map of the wild-type sequence. In either case, our results are better (or equivalent when the statistical signal is too poor) to other DCA-inspired strategies, suggesting that modeling the whole evolutionary trajectory of Directed Evolution experiments allows for a more robust description of the fitness landscape. Moreover, one of the most interesting outcomes of our analysis is the prediction of how experimental strategies could be optimally tuned to produce more informative data. In particular, by running extensive simulations of in-silico experiments, we show how the trade-off between mutation and selective pressure is a critical parameter that needs to be fine-tuned. In agreement with what was observed in [45], our analysis suggests that lower selective pressure in both [26, 29] would have been beneficial to explore more efficaciously the fitness landscape.

## II. NEW APPROACH

*Annealed Mutational approximated Landscape* (AMaLa) uses the sequencing samples of rounds of Directed Evolution experiments to learn a map between the protein amino acid sequence and the fitness associated with the selection process, generically indicated as fitness landscape. Typically, fitness in these experiments is related to the binding affinity to a certain target, or to more complex phenotypic traits, such as antibiotic resistance in bacterial strains.

We consider the probability to observe a generic sequence at a certain time (or round) *t* in the following form:

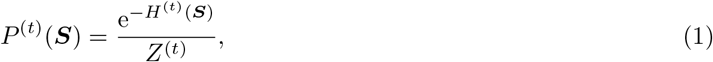

where *H*^(*t*)^(***S***) is a time-dependent Hamiltonian function, and *Z*^(*t*)^ = ∑ _{***S***}_ exp [*−H*^(*t*)^(***S***)] is the associated partition function. The approximation assumes that the model probabilities of observing a given sequence at different times encoded in (1) are statistically independent. An alternative approach would consider the dynamics as a Markov process, describing the probability of whole trajectories in sequence space. However, such a strategy seems to be computationally intractable, as one should sum over all possible trajectories connecting two sequences at subsequent times. Here, we consider a factorized time dependent likelihood, that effectively takes into account the Directed Evolution dynamics, as:

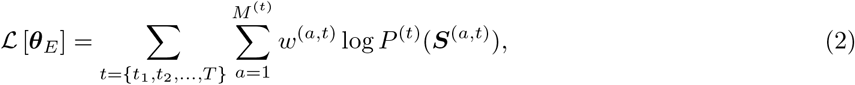

where *w*^(*a,t*)^ is the normalized abundance of sequence *a* = 1, …, *M* ^(*t*)^ at time 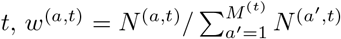, and *N* ^(*a,t*)^ is the absolute abundance of sequence *a* at round *t*. We will implement a maximum likelihood estimate of ***θ***_*E*_, which estimates the model parameters maximizing Eq. (2), the likelihood of the model parameters given the data. The second important assumption is that the time-dependent Hamiltonian function depends on two terms that account for the two different processes occurring in Directed Evolution experiments: selection, and mutation.

To describe the selection term, we introduce a statistical energy *E*(***S***) function of the sequence, labeled by ***S*** = (*σ*_1_, *σ*_2_, …, *σ*_*L*_), where *L* is the number of sites in the sequence, and *σ*_*i*_ = {A, C, …, Y} is the amino acid at residue *i*. We hypothesize that the statistical energy (or more precisely, its opposite) is related to the fitness of the sequence in a selection process. In analogy with standard DCA analysis, we choose to parameterize the energy function as a generalized Potts model:

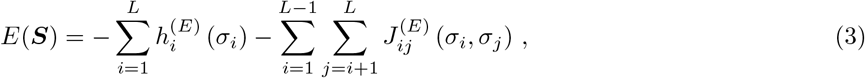

The set of parameters ***θ***_*E*_ := {***h***^(*E*)^, ***J*** ^(*E*)^} is related to the single-site residue frequency and pairs epistatic interactions [52]. The amino acids are mapped onto natural numbers *σ*_*i*_ *∈* 1, …, *q*} where *q* = 20 and *i ∈* {1, …, *L* identifies the sites along the sequence.

Concerning the random mutation process, we used a simple generalization of the Jukes-Cantor model [53] to account for amino acids substitutions instead of DNA base pairs. We introduce this approximation of the real mutation process that does not consider codon bias for simplicity, although the same strategy could be utilized in a more general context such as, for instance, considering a probability transition matrix between codons [45]. Jukes-Cantor is a Markov model which assumes equally probable mutations among amino acids, and it is solely defined by a mutation rate *µ*, through the following transition matrix:

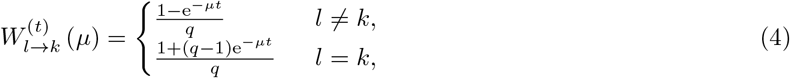

where {*l, k*} 1, …, *q* are two generic amino acids. Eq. (4) defines the transition probability from amino acid *l* to *k* over a time interval *t*. In Directed Evolution experiments, the starting point is typically a single sequence, the *wild-ype*, that undergoes several rounds of error-prone PCR to create the initial library to be screened. Under the hypothesis that mutation is a site independent process, we can express the probability of observing a sequence ***S*** at time *t* through eq. (4) as a function of 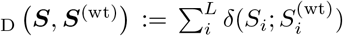, i.e. its Hamming distance from the wild-type, through:

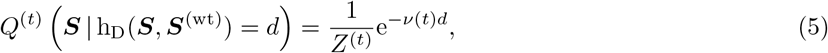

where we introduced the time dependent *v* parameter:

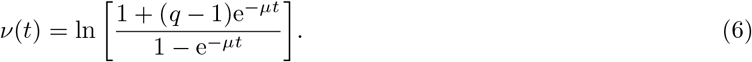

The normalization factor is *Z*^(*t*)^ = e^*−v*(*t*)*L*^ [(*q* 1) + e^*v*(*t*)^] *^L^*. The asymptotic properties of the parameters *v* are the following: As *t →*0 *v* diverges, in such a way that at *t* = 0 all sequences but the wild-type have zero probability. On the other hand, *v* goes to zero as *t →*+ *∞*, so that the asymptotic distribution is uniform in the sequence space. Note that the presented model is defined for simplicity over a continuous-time domain while the real rounds occur at discrete times. Validation on simulated data is analyzed in subsection III B, whereas further results on the purely mutational process (in absence of selection) are reported in the Supplementary Materials section II.

Combining the Jukes-Cantor mutational model with the selection process term, we obtain the Hamiltonian expression and thus the probability of finding a sequence ***S*** at time *t* as:

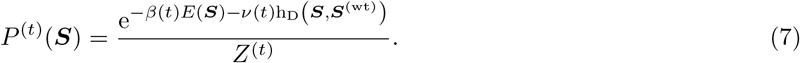

Comparing eq. (7) to the case with only purely random mutation, the presence of the selection biases the variant statistics towards the fittest. As a consequence, in this regime the Jukes-Cantor model becomes at most a convenient approximation. To alleviate these effects, we optimize the parameter *v*(*t*) in eq. (6) to obtain an effective value that maximizes the pseudo-likelihood. When using eq. (7), one assumes that in the course of the Directed Evolution experiment, the consensus sequence does not drift too far away from the wild-type initial sequence. Interestingly, it turns out that in the concrete Directed Evolution experiments considered here this assumption is approximately true.

The *β*(*t*) factor in the selection term encodes its time dependency. From the standpoint of statistical mechanics, it can be interpreted as a fictitious inverse temperature that increases with time, which inspires the term “annealing” in our method name. Let us consider an experiment in the absence of mutation steps (e.g. in the Deep Mutational Scan experiments) but in presence of several selection rounds. We define *P*_*S*_(***S***) as the probability that a sequence ***S*** is selected. The probability *P* ^(*t*)^(***S***) of observing a sequence ***S*** at round *t* is proportional to *P*_*S*_(***S***)^*t*^. Indeed, sequence ***S*** must survive *t* rounds of selection to be observed at round *t*. In this simple case, the inverse temperature *β*(*t*) exactly coincides with *t*. Temperature decreases with subsequent rounds, and in the theoretical limit of an infinite number of selection rounds, the only surviving sequence is the ground state of the Hamiltonian *E, i*.*e*. the sequence with the larger probability according to the model.

The likelihood of the whole experiment outcome is obtained by substituting eq. (7) into eq. (2). As typical in this type of inference problems, the exact maximization of the likelihood requires the determination of the partition function of the model, whose computational complexity scales as 𝒪 (*q*^*L*^). To overcome this limitation, instead of using the likelihood, we maximizes a different but related quantity, called pseudo-likelihood. This approximation allows for a computationally efficient way to learn the parameters [54, 55]. See the supplementary materials section I for the complete definition of the pseudo-likelihood function and the regularization term.

## III. RESULTS

### A. Directed Evolution experiments

We tested AMaLa on three recently published Directed Evolution experiments: two are described in [29] and one in [26]. The proteins mutated and selected in these experiments belong to the *β*-lactamase family (PSE-1 and TEM-1) and acetyltransferase family (AAC6). The *β*-lactamase is responsible for the hydrolysis of antibiotics such as penicillin, ampicillin and carbenicillin while the acetyltransferase is responsible for the catalysis of kanamycin via acetylation. The experiment alternate rounds of variants selection and mutagenesis steps where part of the population is randomly mutated through error-prone PCR. The fitness selection is obtained by exposing bacterial cultures containing the plasmids library to a certain concentration of ampicillin in the case of PSE-1 and TEM-1 (fixed for the former and variable for the latter), and kanamycin for AAC6. In all three experiments, only a subset of the rounds after the selection step are sequenced (see IV for more details on the experimental pipeline).

We used two strategies to test the inferred fitness landscape: (i) by direct comparison of the predicted fitness with experimental measures of the phenotype under selection in the Directed Evolution experiment for a set of variants; (ii) through indirect assessment of the predicted 3D structure of the protein using the inferred epistatic interaction of the learned model (DCA analysis [52]). The first strategy can be applied only to TEM-1, since, to the best of our knowledge, there are no published high-throughput measures of kanamycin and ampicillin resistance for the other two proteins (AAC6, PSE-1). Moreover, being able to use Directed Evolution experiments to predict the structure of a protein is clearly an interesting research perspective in itself, and the main goal of both [29] and [26].

#### Prediction of mutation effect on fitness

High-throughput measurements ampicillin resistance (viz. the same phenotypic trait under selective pressure in [26]) of single site mutants of TEM-1 are presented in [7], whereas measures of minimum inhibitory concentration to *β*-lactamase amoxicillin are presented in [4]. The fitness of the different variants is estimated as the minimum inhibitory concentration concentration to ampicillin of the mutants with respect to the wild-type. It has to be noted that the wild-type sequence in the experiment of [26] (PDB entry 1ZG4) and the one in [7] (Uniprot-P62593) have two mismatches.

The statistical energy score inferred by AMaLa on the dataset of Fantini et al. highly correlates with the Firnberg et al. fitness measurements, with a Pearson correlation coefficient larger than *ρ* = 0.8, suggesting that the method is able to learn a reliable fitness landscape. It is interesting to compare it with the approach outlined in [56], where a Boltzmann learning DCA-based approach is applied to the PFAM *β*-lactamase family (PF13354). In this case, the correlation of the experimental minimum inhibitory concentration with the statistical energy score shows Pearson correlation coefficient *ρ ∼*0.7, as shown in figure 1.

**Figure 1.**
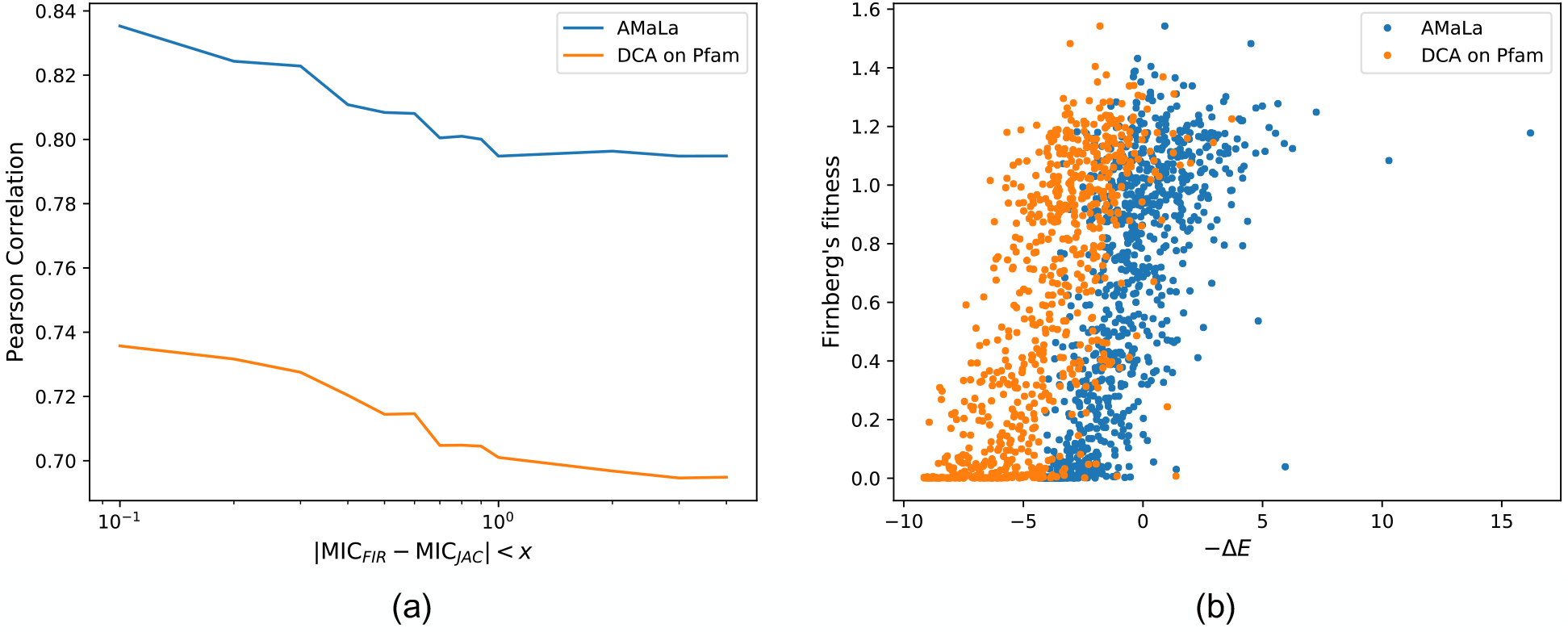
Correlation between inferred energies and fitness measurements realized in [7]. Such measurements are preliminarily mapped over [4] data, following the same procedure proposed in [56]. In this way, it is also possible to filter out nosiest data, retaining only those measurements displaying a low discrepancy between the two datasets. Panel (a) shows the trend of the Pearson correlation obtained as a function of this discrepancy threshold. Namely, correlations are referred to energies inferred over Fantini’s dataset [26] via AMaLa (blue line), and over PFAM PF13354 via PlmDCA (orange). More specifically, such energies are previously mapped over fitness scores via the same procedure exploited to map [7] into [4]. This strategy allows to express the correlation performance in terms of a linear estimator rather than the more general Spearman coefficient. From the plot, it emerges how correlations increase by progressively excluding those measurements with the highest discrepancy among the datasets. Moreover, [26] measurements analyzed via AMaLa turns out to provide a best fitness estimator with respect to the homology family, characterized by a much more dispersed distribution of sequences. On panel (b) the scatter between minus the energies (not mapped), and the fitness measurements of [7] is reported, for a discrepancy threshold between minimum inhibitory concentrations equal to *x* = 1.0.

Directed Evolution experiments from Fantini et al. and the multiple sequence alignment homologous sequences contained in PF13354 provide us with two very different datasets: the first is one is a *local* exploration around the wild-type, with sequences selected to medium-low level of ampicillin selective pressure (average sequence identity of 85%), whereas the second, not surprisingly considering the extremely long time-scale involved in the evolutionary process, shows a remarkably high degree of variability (average sequence identity of 19%). Both can be used to learn a statistical model (AMaLa for [26], PlmDCA [55] for PF13354) providing two distinct sets of model parameters that, remarkably, correlate with each other in terms of statistical energy score (see panel (a) of figure 2), and to the fitness measurements. Interestingly, the parameters of the two models do not correlate with each other (see panels (b)) in figure 2) and consequently, they provide very different contact predictions when used to infer structure information as outlined in the next section. We do not have a clear interpretation of this intriguing result.

**Figure 2.**
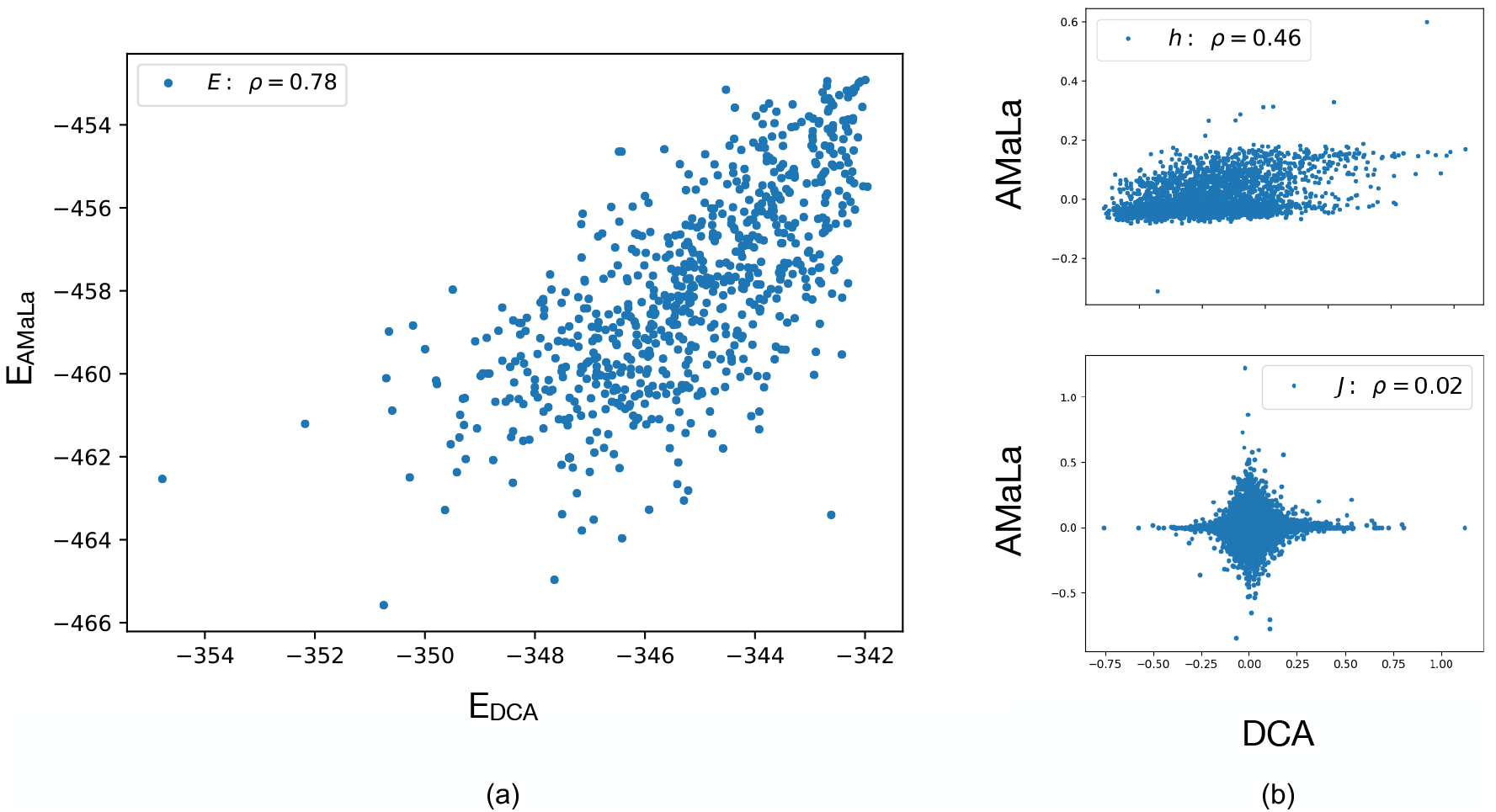
Model comparison between standard PlmDCA performed over the homology family (PF13354) and AMaLa, inferred over [26] data restricted to residues corresponding to the homologues alignment. Panel (a) shows the scatter between the resulting total energies. Panel (b) displays the scatter plots of individual parameters. In the upper plot is reported the scatter among single site fields ***h***, and in the lower the one among pair interaction coupling ***J*** . Even if energetic parameters displays separately either low (*ρ*_*h*_ = 0.46), or no correlation at all (*ρ*_*J*_ = 0.01), the resulting energies are nonetheless significantly correlated, the Pearson coefficient being *ρE* = 0.77. This underlines how the quantity encoding the relevant phenotypic information is indeed the total energy.

#### Residue contacts predictions

Direct coupling analysis is a powerful tool to extract structural properties from multiple sequence alignments of evolutionary-related protein sequences. However, to show its full potential, multiple sequence alignments of at least 10^3^ sequences must be used. For many protein families, the number of homologous sequences available from public databases (e.g. PFAM or UNIPROT) is not sufficient to obtain a reliable folding structure using DCA predictions. Thus, the question of whether one can use artificially created sequences from Directed Evolution experiments to extract structural information, has a very interesting practical purpose as discussed in [29] and [26]. In both papers, the authors apply two similar pseudo-likelihood based inference strategies (the PlmDCA algorithm in [26] and EV-coupling algorithms in [29]) to learn a Potts model from the sequences in the last sequenced round of the experiment. Only one of the two experimental work ([29]) reach a precision sufficient to fold correctly the protein.

Here, we propose a different approach that leverages the sequencing information from all rounds of the Directed Evolution experiment. AMaLa, instead of focusing only on the final step of the in-vitro Darwinian dynamics of Directed Evolution experiments, indeed utilizes the whole time series. We hypothesize that being able to analyze all available data (as opposite to the use of just the last sequenced step) through a model that explicitly (albeit in an effective way) takes into account both mutation and selection steps, could in principle generate a more accurate model of the selection process, providing at the same time better structural information.

We assessed the quality of the DCA scores derived from AMaLa and PlmDCA (the inference method used both in [29] and [26]) by comparing the predicted contact map with true one obtained by PDB structure of the protein (see Material and Methods). The results are shown in figure 3: From the sensitivity plots, we see that, independently from the inference strategy, the predictions for PSE-1 are more accurate than the ones for AAC6. However, if we concentrate on the AAC6 case, AMaLa predictions turn out to be more accurate. As the study of controlled artificial datasets presented seem to indicate, we expect AMaLa to provide better results with respect to PlmDCA when two conditions occur: (i) selection has a relatively weak effect compared to mutation, (ii) not too many rounds of the experiment are performed, so that the Jukes-Cantor modeling of the mutation process remains a good approximation (see next section on in-silico data). The first of the two conditions certainly holds for both proteins, since antibiotic concentration is slightly above the minimum inhibitory one (6*µ*g/mlfor PSE-1 and 10*µ*g/ml for AAC6), while mutation rates are approximately the same for the two. Consequently, since the PSE-1 experiment takes place over 20 rounds, while AAC6 just over 8, we expect to obtain better results in comparison with PlmDCA for the latter rather than the first. Nonetheless, the results obtained on PSE-1 show comparable precision of the two approaches, with AMaLa correctly predicting a higher number of contacts before the first error. Besides, looking at the predicted contact map, the contacts predicted by PlmDCA are mainly close to the polypeptide backbone, while the AMaLa ones are spread over all the contact maps, providing long-range predictions that are more important for constrained molecular-dynamics simulations.

**Figure 3.**
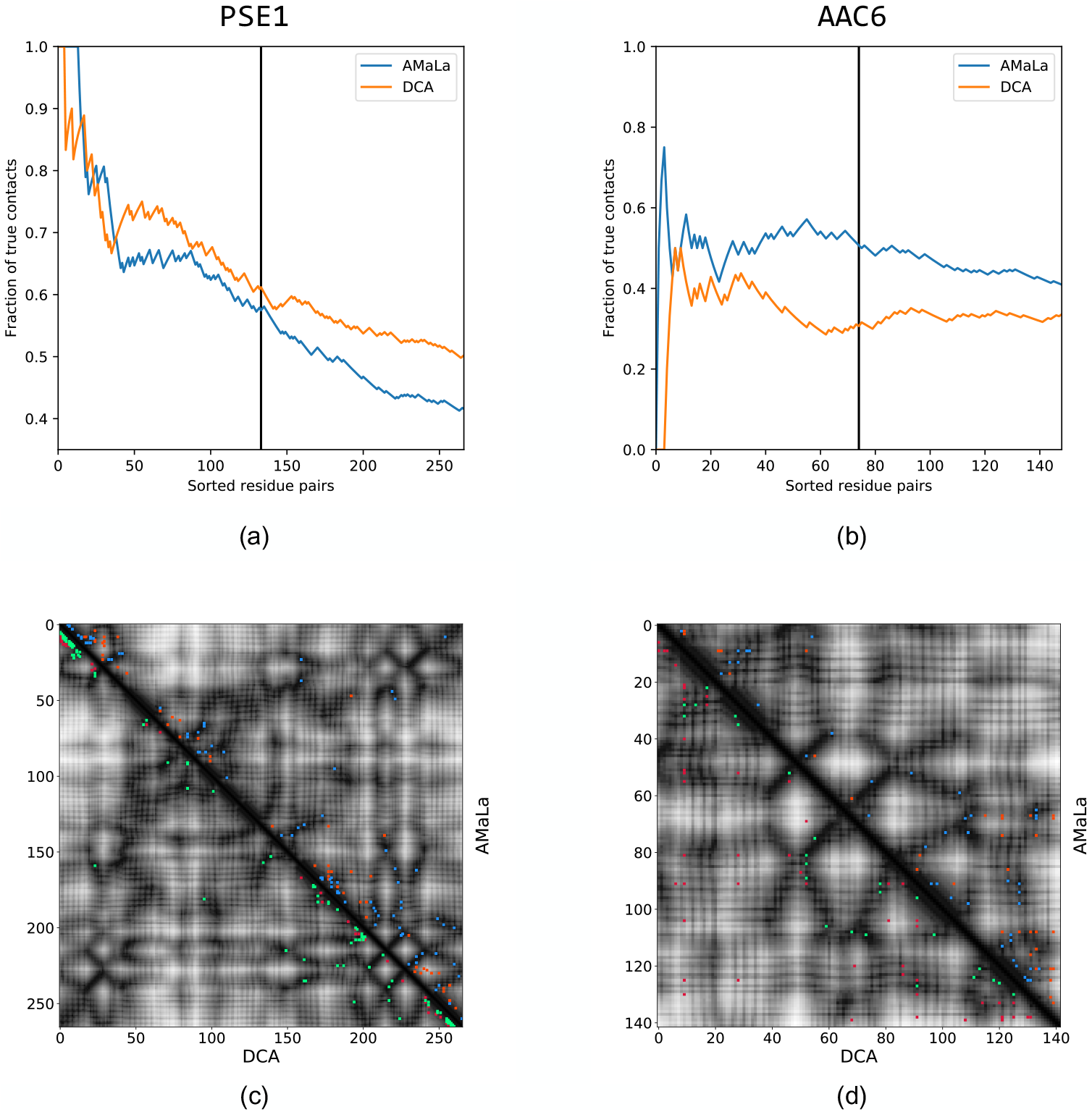
Top: sensitivity plot for contact prediction, via parameters inferred on PSE-1 and AAC6 [29] datasets. Blue curve: the score is computed as the Frobenius norm of the couplings inferred with AMaLa method. Orange curve: the score is computed as the Frobenius norm of the couplings inferred with standard pseudo-likelihood maximization approach. On panel (a) we have the result for PSE-1. At the *L/*2-th ranked residue pair AMaLa provides AUC(*L/*2) = 0.71, PPV(*L/*2) = 0.58, whereas PlmDCA yields AUC(*L/*2) = 0.72, PPV(*L/*2) = 0.61. Panel (b) shows the sensitivity plot for AAC6. In this case AMaLa yields at half of the length AUC(*L/*2) = 0.51, PPV(*L/*2) = 0.51, whereas for PlmDCA we have AUC(*L/*2) = 0.34, PPV(*L/*2) = 0.31. Bottom: contact maps up to *L/*2 predictions. In the upper-right half is reported the results related to AMaLa, whereas in the lower-left is the prediction provided by PlmDCA. Correctly predicted contacts are colored in green/blu, while wrong prediction are reported in red/orange for PlmDCA/AMaLa respectively. Panel (c) reports the result for PSE-1. Even if DCA provides both higher AUC and PPV, AMaLa seems to predict more long range contacts. A similar outcome, although less pronounced, can be appreciated in the panel (d), which shows the contact map related to AAC6.

In complete analogy with what already observed in [26], when the same approach is used for TEM-1 dataset of Fantini et al., neither model is able to provide statistically relevant contact predictions (see supplementary materials section IV). The reason can be related to the different choice of the trade-off between selection strength and mutation rate compared to Stiffler et al. as pointed out in [45]. It is remarkable that, while the model predicts correctly the fitness direct measurements as shown in figure 2, it fails at providing structural information.

Interestingly, in [29] the authors report that the ep-PCR introduces approximately 3-4% amino acid substitutions per round from which we can estimate a mutation rate of *p*_true_ ≃ 0.035. We can compare it with the maximum-likelihood values inferred by AMaLa, that are *p*_infer_ = 0.05 for PSE-1, *p*_infer_ = 0.055 for AAC6, both comparable with the experimentally estimated one.

As a further check, we decided to employ PlmDCA to infer the energy landscape not on the last round only, but on all the available ones. Results in term of contact prediction on [29] data are reported in supplementary materials section IV table III. From the reported values it emerges how extending PlmDCA inference over all sequenced rounds does not provide any significant advantage, but rather seems to produce worse result.

### B. In-silico Directed Evolution experiments

The results on AAC6, PSE-1, and TEM-1 clearly indicate how different experimental conditions (in particular the choice of the mutation rate, and the selective pressure) impact on the ability of the inference algorithm to predict either functional and structural properties. In particular, the interplay between the mutation rate and the selective pressure determines the different dynamical regimes where the assumptions at the basis of our inference method could be more or less verified. To understand the limits of AMaLa, we simulated in-silico Directed Evolution experiments at different ranges of selection and mutation rates (see details in section IV).

The two main parameters of the simulated data are: (i) the site mutation probability parameter *p*, (ii) the strength of the selective pressure 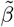: increasing it the selective pressure increases.

In all experiments we keep fixed the teacher energy parameters, and the initial *wild-type* sequence (or more precisely the ground truth energy of such sequence). We used subset of variable size among the total of simulated rounds (typically including between 2 and 5 rounds). The performance of the inference is assessed in terms of the correlation between teacher and student energies computed over a test set of sequences not used to train the model.

In figure 4 we display the retrieval of the true fitness as a function of the mutation rate (panel (a)) and selective pressure (panel (b)). In both cases we observe the existence of an optimal value for both tuned parameters pressure. Interestingly, above the optimal mutation rate, the correlation tends to flatten at a value which is not far from the optimal one, ensuring that AMaLa sweet spot for inference (at fixed selective pressure) is in general towards a high mutation rate regime. Just as a reference to real Directed Evolution experiments, the mutation rate reported in [29] is *p*_true_ ≃ 0.035.

**Figure 4.**
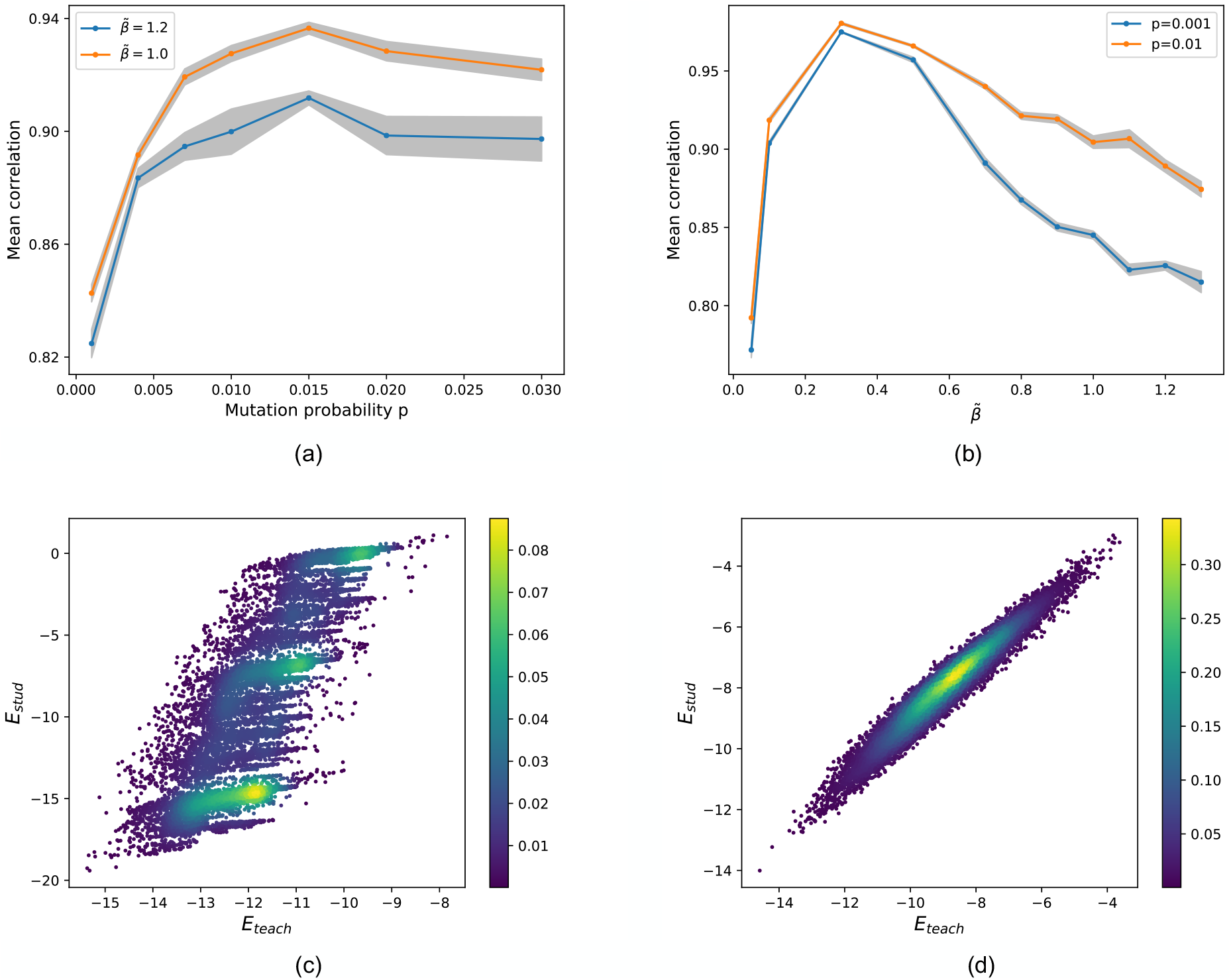
Simulated experiments varying the mutation rate and the selective pressure. On the top panels are shown the Pearson correlation between true and predicted fitness or equivalently teacher and student model energy. In order to estimate statistical fluctuation on the correlation, for each point several experiment replica has been realized (*N*sim = 20-40), reporting mean and standard deviation. On panel (a) it is varied the mutation rate for two choices of the selective pressure: 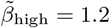 (blue), 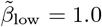 (orange). Conversely on panel (b) it is varied the selective pressure at two fixed mutation rates: *p*_low_ = 0.001(blue), *p*_high_ = 0.01(orange). An optimal mutation rate seems to emerge, with the mean Pearson coefficient which flattens for higher mutations rate. Again, performances appear to decrease with increasing selective pressure. Moreover, the curve coinciding with *p*_high_ displays significantly higher correlations. The bottom panels show two examples of density scatter plot between true (*x*-axis) and inferred (*y*-axis) energies over the test set. Two limiting cases are shown: high selective pressure and low mutation rate in panel (c) (*p* = 0.001 and 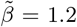), and low selective pressure and high mutation rate in panel (d) (*p* = 0.05 and 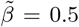) where AMaLa recover the right fitness landscape. In the former case the Pearson’s correlation is *ρ* = 0.81, while in the latter is *ρ* = 0.97.

Unfortunately, we do not have access experimentally to a quantitative assessment of the strength of the selective pressure, making a direct comparison with experiments difficult. The method works at intermediate selective pressure as the selection tend to undermine the method assumptions (see section II). Indeed, when the selection strength is too low (depending on the time scale of the experiment), the sequence dynamics is dominated by genetic drift and, not surprisingly, the correlation between teacher and student degrades. The degradation of the performance observed for higher selective pressure is due to a combination of effects: on the one hand, we expect the in the limit of *β → ∞* only the lowest energy sequence generated in the mutation step would survive, making any inference impossible. On the other hand, at intermediate but high selective pressure, we expect that the consensus sequence starts drifting significantly from the initial wild-type sequence, making the drift term of eq. (7) an inaccurate description of the purely mutational step.

In Directed Evolution experiments, one of the limiting factor is the number of selection rounds that can be sequenced (and that therefore can be used for the inference). In the following we will assume we can afford only between two and five rounds of sequencing, and we ask which rounds bring the larger information content.

As shown in figure 5 panel (a), for PlmDCA the correlation between the teacher and student energies of the test set increases as a function of the last round time, whereas AMaLa performance behaves just in the opposite way: earlier round times give better results. This finding is particularly interesting as it suggests that by using AMaLa one could achieve better inference results by performing just a limited number of rounds, i.e. with a lower experimental effort. However, AMaLa overall performance are always better then PlmDCA for any choice of the sequencing round.

**Figure 5.**
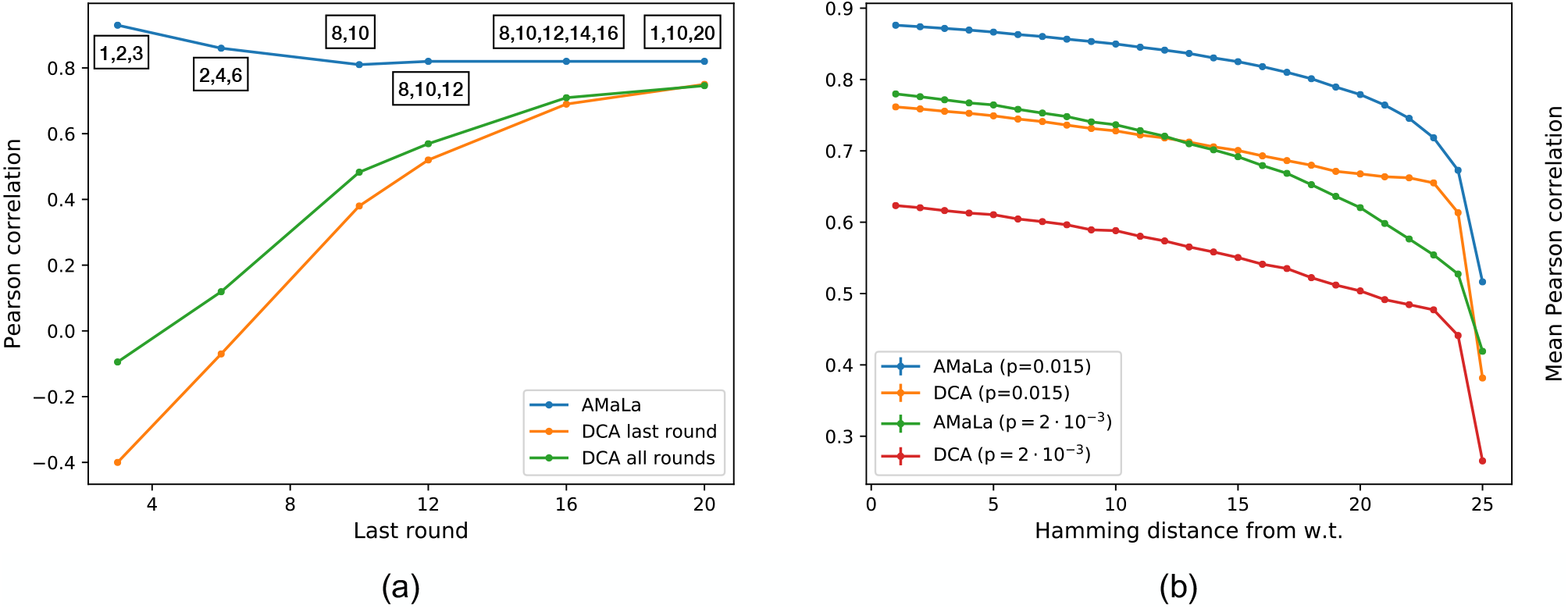
Inferred signal dependence on the number of rounds and hamming distance from wild-type. Left panel (a): Pearson correlation when the number of rounds are changing. Comparison between PlmDCA on the last round (or all rounds) and AMaLa. The sequenced rounds are: (1, 2, 3); (2, 4, 6); (8, 10); (8, 10, 12); (8, 10, 12, 14, 16); (1, 10, 20). PlmDCA significantly depends on the number of performed rounds, not significantly inferring the fitness landscape up to round *∼*16. On the contrary, AMaLa provides predicted energy functions highly correlated with the fitness even for low number of performed selection rounds. Right panel (b): Degradation of mean Pearson correlation between inferred and true energies, as a function of the Hamming distance from the wild-type sequence. Two different simulation are considered: high (*p* = 0.015) and low (*p* = 0.002) mutation probability. Changing such parameter varies the broadness of the library screening during an experiment, resulting in probing a more local or more broad region of the sequence space. AMaLa predictions are systematically better than PlmDCA, while the latter display a slower decrease in correlation augmenting the distance from wild-type.

Furthermore, the in-silico experiments can be used to investigate the generalization power of the learned fitness landscape beyond the local region of sequence space probed by the experiment. More specifically, how far from the wild-type an inference strategy is still able to predict the fitness? To answer to this question we trained both AMaLa and PlmDCA on rounds (2,4,8). Then, we tested the teacher student energy correlation over randomly extracted sequences at Hamming distance up to the whole sequence length (here *L* = 25). As shown in figure 5 panel (b), we can see that that both in the case of low and high mutation rate: (i) over the whole range of hamming distance from the wild-type sequence, AMaLa always shows higher correlation with the teacher energies, (ii) PlmDCA performance seem to degrade more slowly as a function of the distance from the wild-type sequence.

## IV. MATERIAL AND METHODS

### Experimental Pipeline

The experiments involve repeated rounds of mutation and selection, starting from a natural sequence, named *wild-type*. Repeated cycles of error-prone PCR are applied to the library at each round to introduce mutations. Functional selection is obtained by inserting the plasmids with the variants in a bacterial colony and then placing the colony in an environment with a relatively low antibiotic concentration: 6 *µ*g/mL ampicillin for PSE-1, 10 *µ*g/mL kanamycin for AAC6, whereas two different ampicillin concentrations are used for TEM-1, namely 25 *µ*g/mL for all rounds but 5 and 12, when the concentration is raised to 100*µ*g/mL. For a subset of the rounds (with the last round included), a sample of the population after the selection is sequenced. Thus, for each sequenced round *t* we obtain the abundances *N* ^(*a,t*)^ for the variants *a* = 1 …, *M* ^(*t*)^ with a.a. sequence *S*^(*a,t*)^, *M* ^(*t*)^ being the number of unique sequence present in the sample at time *t* (see the figure 6 for the full pipeline).

**Figure 6.**
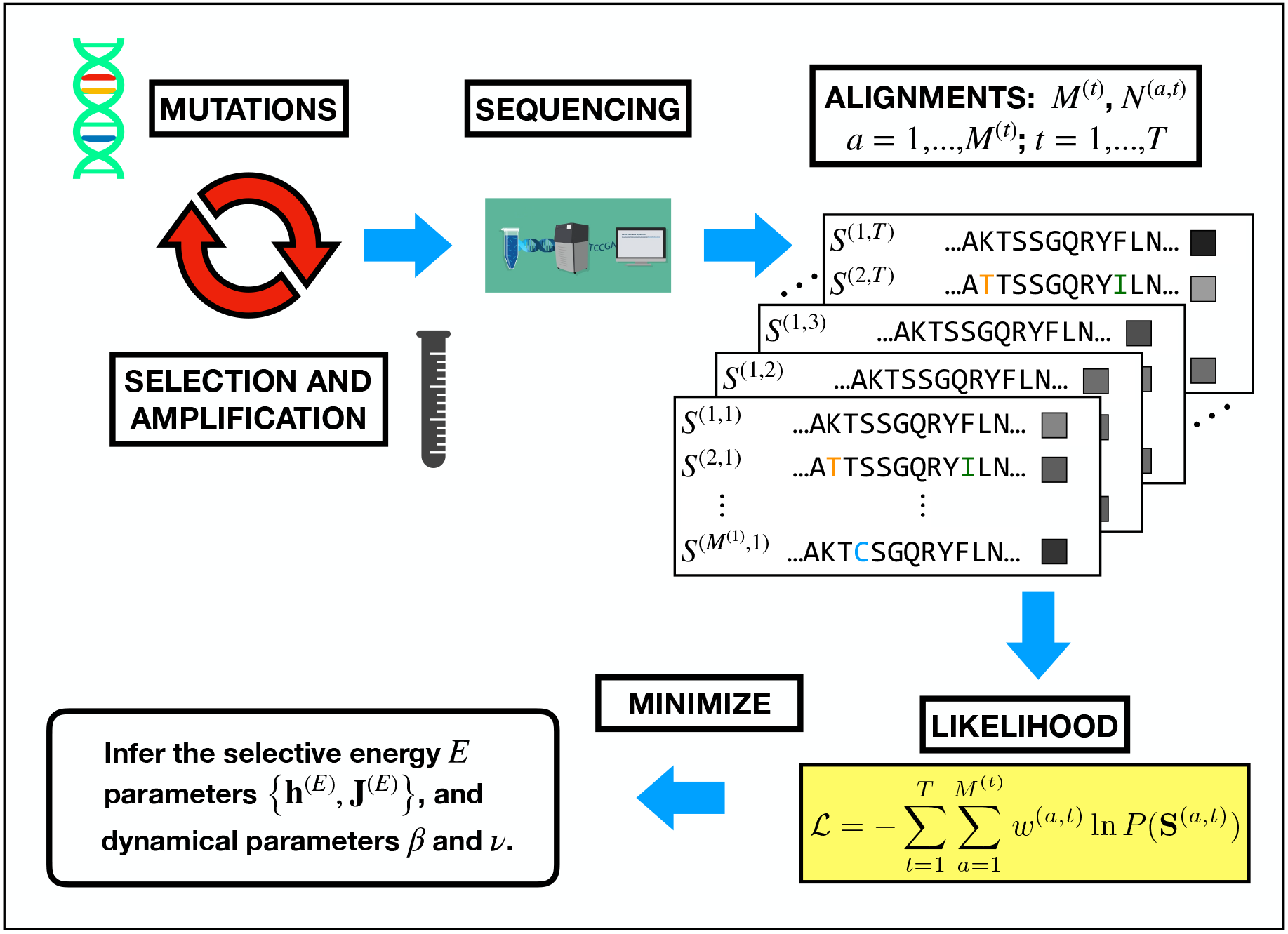
Pictorial representation of data generation in a Directed Evolution experiment and how they are plugged into the likelihood function to perform the inference. The sequencing of repeated rounds of mutation and selection generates a set of multiple sequence alignments. There, we highlighted with colored letters the sites which have been mutated with respect to the wild-type, coinciding with the first row of the alignment. Moreover, at the right of the sequences, boxes in gray-scale represents the abundances (increasing from black to white). Each sample of sequences *{****S***^(*t*)^*}* and the related abundances *{****N*** ^(*t*)^*}*, for *t* = 1, …, *T* are used to define the likelihood function, which subsequently depends only on the parameters to be inferred: ***θ****H* = {***h***^(*E*)^, ***J*** ^(*E*)^, ***v, β}*** (see eqs. (3),(1),(2) and (7)). The inference of these parameters is based on the maximization of the log-likelihood. In order to determine the parameters ***β*** and ***v***, the maximization problem over the energetic parameters is repeated, performing a scan over a set of possible values. Then the pair (***β***opt, ***v***opt) corresponding to the global maximum of the minus log-likelihood is retained.

### Model learning

The abundances of the sequenced variants are used to compute the normalized weights *w*^(*a,t*)^ in (2). To learn the parameters (***v, β, θ***_*E*_) of the model (7) we maximize (2) using a pseudo-likelihood approximation (see supplementary materials section I). Although (2) is convex with respect to the energy parameters ***θ***_*E*_ and the inverse temperatures ***β*** separately, it is no longer true when the parameters are varied simultaneously. Two strategies are possible: (i) optimizing the energetic parameters at different values of ***β*** and select the maximal pseudo-likelihood. (ii) Starting from an arbitrary ***β*** and optimize sequentially ***θ***_*E*_ and ***β*** using gradient descent algorithm (e.g. Newton algorithm). Since the ***θ***_*E*_ is defined up to a multiplicative constant, we can set *β*(*t*_1_) = 1 (with *t*_1_ the time of the first sequenced round) or alternative *β*(*t* = *T*) = 1 without losing generality. The last choice is mandatory in the case in which *β*(*T*) diverges. In such a scenario *β*(*t*_1_) *≡*0 and the intermediate values are constrained in the domain [0, 1]. In order to set an optimal value for the Jukes-Cantor parameter *v*(*t*), we decided to maintain the functional form in eq. (6), thus performing a scan over the possible values of the mutation rate *µ*. The value 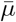 yielding a maximum for the pseudo-likelihood then defines each component for the different round time: 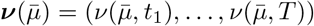.

### Contacts prediction

To compute an epistatic score associated with each pair of residues we used the Frobenius norm with APC correction (originally introduced in [57]. For each pair of positions *i* and *j* the Frobenius norm over all possible amino acid combinations of the interaction parameters *J*_*ij*_ is computed:

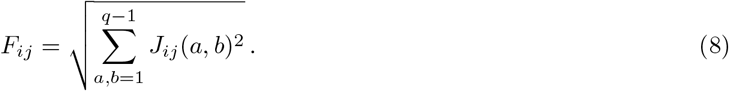

As the Frobenius norm is not-gauge invariant, is it important to transform first the Potts parameters in zero-sum-gauge [55]. Lastly, we applied the average product correction (APC) to the *F* matrix: 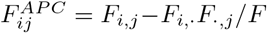_*·,·*_, where the dot represents the average over the index. The same procedure has been used to obtain both AMaLa and PlmDCA derived DCA scores.

To assess the predicted contact maps we compared it to the residues contact extracted from crystal structures present in the PDB database (1G68 for PSE-1, 4EVY for AAC6 and 1ZG4 for TEM-1). Two residues are in contact if at least two heavy atoms have a distance less than 8°A. We consider only residues with a separation on sequence |*i − j*| *≥* 5.

### Experiment simulation

To simulate Directed Evolution experiment we define a dynamical process that mimics the mutation and selection steps occurring in a real experiment. We define *N* ^(*a,t*)^ as the number of clones of variant *a* present at the round *t* for *t ∈ {*1, …, *T}*. The total number of clones is kept fixed along the simulation and equal to 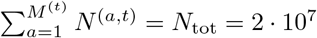.

Mutations are drawn from a site independent uniform distribution over the space of 20 amino acids. The unique parameters we consider is the mutation probability *p* (or equivalently the mutation rate *µ* see supplementary materials section II). For every clone of a given variant, the number of sites to be mutated is drawn from a binomial distribution of probability *p*.

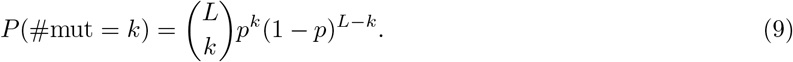

In practice, for each selected site the new mutations are extracted uniformly over the possible different amino acids. This process either generates new variants or increases the abundances of already present ones.

Finally, we simulate the selection step by associating a *survival* probability *P*_*S*_(***S***^(*a*)^) to each variant *a* via a Boltzmann weight proportional to 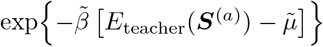.

The energy function *E*_teacher_ has the same functional form of equation (3) and given its parameters it constitutes the ground-truth fitness landscape (the *teacher* model). The parameter 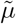 is a sequence independent chemical potential that fixes the scale of the binding probability. Typically numerical values employed in the simulations are around 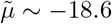.

From the set of variants produced by the mutation process, a subset *n*^(*a,t*)^ of surviving clones is selected according to a binomial process defined by:

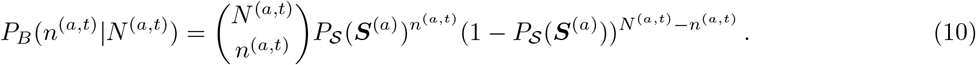

Finally, the population of clones that survived the selection step is amplified up to a fixed number *N*_tot_ according to the following multinomial distribution:

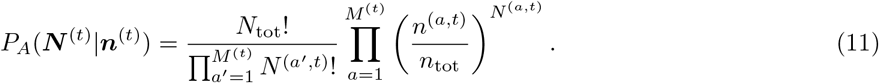

In addition, we randomly sample *R*_tot_ = 10^6^ sequences out of the ***N*** ^(*t*)^ present variants to introduce the sampling noise and simulate the effect of the sequencing.

In figure 1 in supplementary materials, is reported a pictorial representation of the whole pipeline for the generation of simulated data.

The parameters setup of the simulation was chosen with the aim to be as close as possible to a real experiment and to not introduce unnecessary artificial features. The teacher model for the ground truth fitness landscape is obtained by the inference of a Potts model on a Deep Mutational Scan (DMS) experiment [58].The inference method used to obtain the teacher model is described in [51] and it provides a reliable model of the fitness in the absence of mutagenesis steps. In the considered (DMS) experiment the WW domain of the hYAP65 protein has been mutated and selected to bind to its cognate peptide ligand. The mutated part of the protein has a length *L* = 25 amino acids.

While finalizing this work, we became aware of a similar approach described in [45]. Their strategy relies on a simultaneous treatment of selection and mutagenesis. The fitness approximated landscape is inferred over the homologous alignment, specifically via Boltzmann learning of a generalized Potts model. Such energy provides a proxy for fitness, and a tool to probe context dependent mutations, for the energy function includes couplings between different residues. Indeed, a MCMC is implemented to generate a library which mimics the one that would have been obtained in a real Directed Evolution experiment. The elementary step of this MCMC includes both mutation and selection. The energy variation of single site mutations with respect to the wild-type defines the acceptance probability (which depends only on the a.a. sequence). On the other hand, the proposed mutations are restricted to the allowed single mismatch transitions among codons 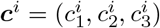, thus involving the genomic sequence. This may suggest a possibility to improve AMaLa itself: the Hamming distance in the Jukes-Cantor contribution in (7) may be computed over the genome alignment. In this away forbidden transitions among a.a.’s are automatically excluded, but at the same time also multiple transitions are allowed, even if exponentially suppressed. Remarkably the findings of [45] with respect to the optimal regime for a Directed Evolution experiment agrees with the results we derived from the application of AMaLa to both in silico and in vitro data.

## V. DISCUSSION

In this work, we presented AMaLa, a new inference strategy for modeling Directed Evolution experiments. At the heart of our algorithm, there is an effective modeling of the two main ingredients of Directed Evolution experiments: mutation and selection. Whereas other competing algorithms’ computational strategies typically use data from the last sequenced round, our model leverages all the available *history* of the experiment in terms of all sequenced rounds of selection. By doing so, we are able to infer a better statistical model both in terms of the ability to predict functional phenotypes and structural properties of the protein.

As Directed Evolution is becoming a very relevant instrument to test on a controlled ground different evolutionary theories, as well as an invaluable tool to find optimal target phenotype sequences with pharmaceutical and/or biotechnological interest, we believe that a reliable statistical modeling of the experiment has a twofold interest: on the one hand, we show how our model can predict quantitatively the trait under selection also of variants that have been not used in the training data, suggesting that our model could in principle be used to propose sequences, and/or libraries of sequences of improved biological activity; on the other hand, our statistical model could be used to optimally set the experimental control parameter. In particular, we were able to stress how relevant is the trade-off between mutation and selection in different experiments. Our findings are also corroborated by extensive in-silico Directed Evolution experiments, where the modification of these two parameters can be easily taken into consideration in limits that would otherwise be experimentally inaccessible. AMaLa of course is a first attempt at modeling evolutionary trajectories under fixed selection pressure, although some interesting attempts have been recently published in [59] in the somehow different context of the inference of genetic linkage in population genetics. In particular, the way in which we model the mutation step could be made more accurate by taking into account codon biases and more realistic transition probabilities (the first attempt in this direction has been proposed in [45]). From this point of view, our work suggests that it might be useful to sequence the library before and after the selection step to disentangle the effect of mutation and selection, and produce a better correlation between statistical energies and the empirically measured trait under selection.

## ACKNOWLEDGMENTS

The authors acknowledge financial support from Marie Sklodowska-Curie (grant agreement no. 734439) (INFERNET). We also wish to thank Martin Weigt and Matteo Bisardi for an enlightening conversation on the trade-off between selection and mutation in Directed Evolution experiments, and Marco Fantini for sharing with us interesting details of his experiments.

