## Supplementary for "AMaLa: Analysis of Directed Evolution Experiments via Annealed Mutational approximated Landscape"

### Supplementary material for: Application of Pseudo-likelihood Inference to Phenotype selective Experiments

Luca Sesta† and Guido Uguzzoni †
*Politecnico di Torino, Corso Duca degli Abruzzi 24, I-10129, Torino-Italy*

Jorge Fernandez-de-Cossio Diaz *
*Laboratory of Physics of the Ecole Normale Supérieure,*
*CNRS UMR 8023 & PSL Research, Paris, France;*
*Systems Biology Department, Center of Molecular Immunology, Havana, Cuba*

Andrea Pagnani ‡
*Italian Institute for Genomic Medicine, IRCCS Candiolo,*
*SP-142, I-10060 Candiolo (TO) - Italy and*
*INFN, Sezione di Torino, Torino, Italy*
(Dated: July 26, 2021)

#### I. PSEUDO-LIKELIHOOD

In this section, we recall briefly the concepts and methodologies defining the pseudo-likelihood inference procedure. For further information we recommend references [1, 2].

Each protein variant is encoded in a sequences of 20 amino acids and eventually an additional symbol for the alignment gap $\boldsymbol{S} = (\sigma_1, ..., \sigma_L)$. We associate to each sequence an energy value given by the following Hamiltonian function (Potts model):

$$H\left(\boldsymbol{S}\right) = -\sum_{i=1}^{L} h_i(\sigma_i) - \sum_{i=1}^{L-1}\sum_{j=i+1}^{L} J_{ij}(\sigma_i, \sigma_j), \tag{1}$$

defined by the set of fields and couplings $\{\boldsymbol{h}, \boldsymbol{J}\}$. The probability of a variant is given by the Boltzmann weight $\mathrm{e}^{-H(\boldsymbol{S})}/Z$, where $Z = \sum_{\{\boldsymbol{S}\}} \exp\left\{\sum_{i=1}^{L} h_i(\sigma_i) + \sum_{i=1}^{L-1}\sum_{j=i+1}^{L} J_{ij}(\sigma_i, \sigma_j)\right\}$ is the partition function.

The log-likelihood associated with a particular parameters choice $\{\boldsymbol{h}, \boldsymbol{J}\}$ given the data sample is

$$\mathcal{L}\left[\boldsymbol{h}, \boldsymbol{J}\right] = \sum_{a=1}^{M} w^{(a)} \log P(\sigma_1^{(a)}, ..., \sigma_L^{(a)}), \tag{2}$$

where $w^{(a)}$ is the empirical frequency of the variant $a$.

In the maximum-likelihood approach the inference problem entails the determination of the parameters $\{\boldsymbol{h}, \boldsymbol{J}\}$ that maximize $\mathcal{L}$. However, this method inevitably entails the computation of the global partition function $Z$, an operation which is computationally unfeasible due to the large size of the phase space, which is $q^L$ (with $q$ either equal to 20 or 21). The necessity to overcome this difficulty leads to the so-called pseudo-likelihood approximation, which maximizes a different function based on single-site conditional probabilities:

$$P\left(\sigma_r = \sigma_r^{(a)} | \sigma_{-r} = \sigma_{-r}^{(a)}\right) = \frac{\exp\left\{h_r(\sigma_r^{(a)}) + \sum_{i\neq r} J_{ri}(\sigma_r^{(a)}, \sigma_i^{(a)})\right\}}{\sum_{l=1}^{q} \exp\left\{h_r(l) + \sum_{i\neq r} J_{ri}(l, \sigma_i^{(a)})\right\}}, \quad a = 1, ..., M. \tag{3}$$

Averaging over the data sample the log of equation (3), yields the single site contributions to pseudo-likelihood:

$$\mathcal{L}_r^{\text{pseudo}} = \sum_{a=1}^{M} w^{(a)} \log\left[P\left(\sigma_r = \sigma_r^{(a)} | \sigma_{-r} = \sigma_{-r}^{(a)}\right)\right], \tag{4}$$

where we labeled as $\boldsymbol{h}_r$ the set $\{h_r(l)\}_{l=1}^{q}$ and $\boldsymbol{J}_r = \{J_{ri}(l,k)\}_{l,k=1}^{q}$ and $i \neq r$. The inference procedure is then based on the minimization of the sum over sites of (4), plus a regularization term, that is: $\mathcal{L}_{\text{pseudo}} + R\left[\boldsymbol{h}, \boldsymbol{J}\right] = \sum_{r=1}^{L} g_r(\boldsymbol{h}_r, \boldsymbol{J}_r)$. The regularization is of type $l_2$, indeed its single site contribution reads:

$$R_r\left[\boldsymbol{h}_r, \boldsymbol{J}_r\right] = \lambda_h \|\boldsymbol{h}_r\|_2^2 + \lambda_J \|\boldsymbol{J}_r\|_2^2 = \lambda_h \sum_{l=1}^{q} h_r(l)^2 + \lambda_J \sum_{i\neq r}\sum_{l,k=1}^{q} J_{ri}(l,k)^2, \tag{5}$$

where we introduced the regularization multipliers for fields and couplings $\lambda_h$ and $\lambda_J$. This kind of regularization is equivalent in a Bayesian framework to assume a gaussian prior over the parameters.

The pseudo-likelihood function is asymptotically equivalent to (2), they provide the same results in the limit of a very large data sample.

This approach has two different advantages: (i) differently from the global partition function $Z$, its single site version $Z_r(a) = \sum_{l=1}^{q} \exp\left\{h_r(l) + \sum_{i\neq r} J_{ri}(l, \sigma_i^{(a)})\right\}$ can be computed. (ii) each $L$ terms related to different sites can be optimized independently, so that computation can be performed in parallel (asymmetric minimization). A drawback of this strategy is that it provides two different coupling estimates $J_{ri}^{r*}$, $J_{ir}^{i*}$ for each pair of sites $r$ and $i$ that can be averaged to obtain a the final estimation $J_{ij} = (J_{ij}^{i*} + J_{ji}^{j*})/2$.

#### II. MODELING THE PURELY MUTATIONAL PROCESS

The purely mutational process, a process without functional selection, is described by a generalized Jukes-Cantor model. The model is defined by eq. (4) in the main text, which expresses the probability of an amino acid change in a time interval $t$. These transition probabilities depend on two parameters: the number of amino acids $q = 20$ and the single site mutation rate $\mu$. The single site mutation probability $p$ is obtained from $\mu$ by setting $t = 1$ in eq. (4) in the main text, and summing over all possible $q - 1$ mutations:

$$p = \frac{q-1}{q}\left(1 - \mathrm{e}^{-\mu}\right). \tag{6}$$

The transitions probability can be combined in order to obtain the probability of observing the sequence $\boldsymbol{S}$ at any time, in the hypothesis of a unique common wild-type sequence:

$$\begin{aligned} Q^{(t)}\left(\mathbf{S}|\mathrm{h_D}(\boldsymbol{S},\boldsymbol{S}^{(WT)}) = d\right) &= \left(\frac{1}{q}\right)^N \left(1-\mathrm{e}^{-\mu t}\right)^d \left(1+(q-1)\mathrm{e}^{-\mu t}\right)^{N-d} \\ &= \left(\frac{1}{q}\right)^N \exp\left\{d\ln\left[1-\mathrm{e}^{-\mu t}\right] + (N-d)\ln\left[1+(q-1)\mathrm{e}^{-\mu t}\right]\right\} \\ &= \left(\frac{1}{q}\right)^N \left[1+(q-1)\mathrm{e}^{-\mu t}\right]^N \exp\left\{-d\left[\ln\left(\frac{1+(q-1)\mathrm{e}^{-\mu t}}{1-\mathrm{e}^{-\mu t}}\right)\right]\right\} \\ &= \frac{1}{Z^{(t)}}\mathrm{e}^{-\nu(t)d}, \end{aligned} \tag{7}$$

where we introduced the time dependent parameter $\nu(t)$. We can express the normalization factor $Z^{(t)}$ as a function of $\nu$ and $q$ only:

$$Z^{(t)} = \sum_{\{\boldsymbol{S}\}} \mathrm{e}^{-\nu(t)\mathrm{h_D}(\boldsymbol{S},\boldsymbol{S}^{(WT)})} = \sum_{d=0}^{L}\binom{L}{d}(q-1)^d\mathrm{e}^{-\nu(t)d} = \mathrm{e}^{-\nu(t)L}\left[(q-1)+\mathrm{e}^{\nu(t)}\right]^L. \tag{8}$$

Finally, we can also perform a change of variable, employing eq. (7) to compute the first two moments of the Hamming distance from the wild-type:

$$\begin{cases} \langle d(t)\rangle = \frac{1}{Z^{(t)}}\sum_{d=1}^{L} d\binom{L}{d}(q-1)^d\mathrm{e}^{-\nu(t)d} = \frac{(q-1)L}{(q-1)+\mathrm{e}^{\nu(t)}}, \\ \langle d^2(t)\rangle - \langle d(t)\rangle^2 = \frac{(q-1)L\mathrm{e}^{\nu(t)}}{\left[(q-1)+\mathrm{e}^{\nu(t)}\right]^2}. \end{cases} \tag{9}$$

#### III. SIMULATED DIRECTED EVOLUTION

The in silico simulation of the directed evolution experiment comprises the following steps. A sequence is chosen to be the wild-type of the simulated experiment. From this unique common origin, new sequences are generated simulating the mutagenesis via a binomial site-independent process, defined by a single site mutation probability $p$. The newly generated sequences are then selected according to an energy function defined by a set of parameters $\left\{\boldsymbol{J}^{(E)}, \boldsymbol{h}^{(E)}\right\}$. More precisely, the selection probability for a sequence $\boldsymbol{S}$ is proportional to the Boltzmann weight $P_{\mathcal{S}}(\boldsymbol{S}) \propto \exp\left[-\tilde{\beta}(E(\boldsymbol{S}) - \tilde{\mu})\right]$, where is $E$ is the generalized Potts model defined by the aforementioned set of fields and couplings. The auxiliary parameters $\tilde{\beta}$ and $\tilde{\mu}$ can be employed to change the shape of the selection probability distribution. The parameter $\tilde{\beta}$ has been chosen to satisfy the *rare biding* condition. The total number of clones is held fixed during the experiment ( $N_{\mathrm{tot}} = 2 \cdot 10^7$). Cycles of mutation and selection (and amplification) are repeated for a total of $T = 20$ rounds. Then, a subsampling process is performed to obtain the sequencing reads, with $R_{\mathrm{tot}} = 1 \cdot 10^6$ total number of reads. Figure 1 shows the pipeline for the generation of in silico data.

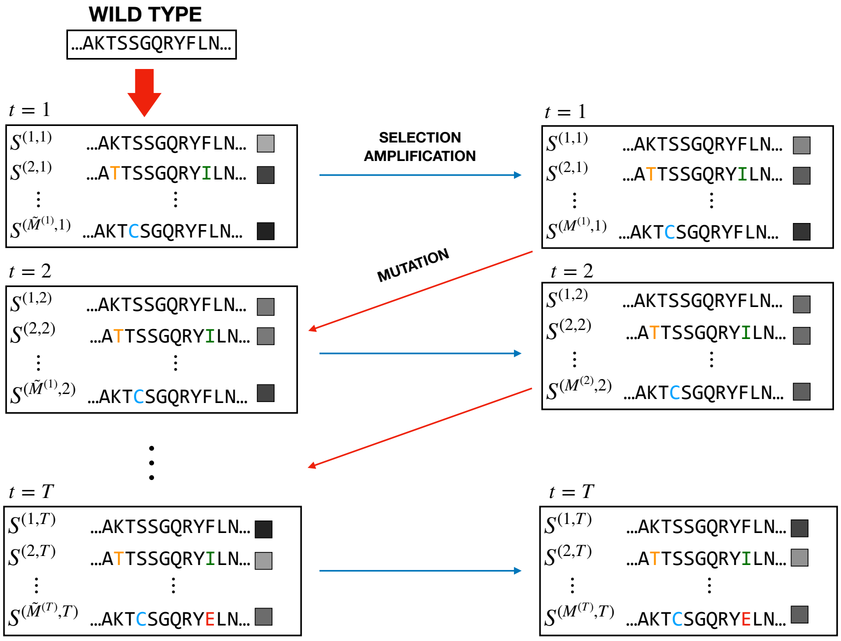


Figure 1. Schematic representation of the pipeline for the generation of in silico data. The starting point is a host of copies of the wild-type sequence. The total abundance $N_{\text{tot}}$ remains constant through the entire simulation. Then, mutagenesis is performed over this collection of wild-types, obtaining a library of $\tilde{M}^{(1)}$ unique sequences. Mutated residues are colored, and the gray-scale boxes indicate the abundances related to each unique sequence (increasing population going from black to white). This library is then subjected to both a selection and an amplification step. As a consequence, the abundances vary according to sequences fitness. Moreover, since the number of unique sequences may change, we have a new library size labeled by $M^{(1)}$. The described steps represent the fundamental unit of the simulation, which is then realized by cycling multiple times this block. As a consequence, we obtain two temporal series of alignments: one which stems from mutagenesis $\{\tilde{\boldsymbol{N}}^{(1)}, \tilde{M}^{(1)}; ...\tilde{\boldsymbol{N}}^{(T)}, \tilde{M}^{(T)}\}$, and the other from selection and amplification $\{\boldsymbol{N}^{(1)}, M^{(1)}; ...\boldsymbol{N}^{(T)}, M^{(T)}\}$. Since experimental libraries are typically sequenced after selection only, we retain just the second series of alignments to make up our data sample. To be more precise, a further subsampling process is performed yielding the reads trajectory $\{\boldsymbol{R}^{(1)}, M^{(1)}; ...\boldsymbol{R}^{(T)}, M^{(T)}\}$. For further information, take a look to the dedicated paragraph in section IV of the main text.

In figure 2 is shown the time evolution of the number of variant during the simulation. Two scenarios are depicted, one where it increases steadily and another where it drops after the first selection round.

To investigate how the exploration of sequence space is affected by the selection process we show in figure 3 the average Hamming distance from the wildtype at each round comparing with a process where only mutations occur, the effect of selection is a mild distortion of the trajectory of the purely mutational case. Moreover, the distance of the consensus sequence to the wildtype gives us the information on how the process is centered on the wildtype sequence that is an assumption of the model. In panel (b) of figure 3 is reported the outcome obtained for $p = 0.01$, the result is strongly affected by the fitness landscape and in particular if the wildtype is locally an optimum.

##### A. Inference on the simulated data

In this section, we report some further analysis on the model learning outcome using in silico generated data.

In figure 4, we compare the teacher and student parameters $\left\{\boldsymbol{h}^{(E)}, \boldsymbol{J}^{(E)}\right\}$ using the dataset of figure 4 panel (c) and (d) in the main text. The *true* and inferred energies have a higher correlation than the parameters, arguably it is due to the degeneracy of the energy function, different parameter's choice giving similar energies. The inference provides a solution on the degenerate region that can have parameter's correlation of $\rho_h = 60$, $\rho_J = 40$ while energy correlation is higher $\rho_E = 0.81$.

The figure 5 concern the determination of the optimal values of $p$ ( and thus $\boldsymbol{\nu}$) and $\boldsymbol{\beta}$ parameters maximizing

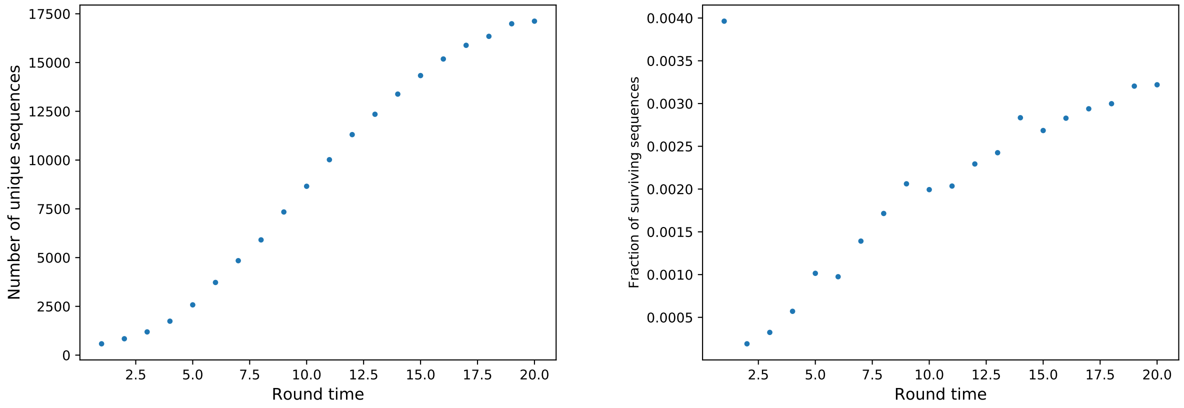


(a) (b)

Figure 2. Evolution of the number of sequences through the experiment simulation. Panel (a) shows the trend of the number of unique sequences, which displays a maximum after 17 rounds. Panel (b) fraction of the newly generated sequences surviving the selection process. Apart for the steep decrease after the first round, this number increases until round 17, when it starts to decrease as well.

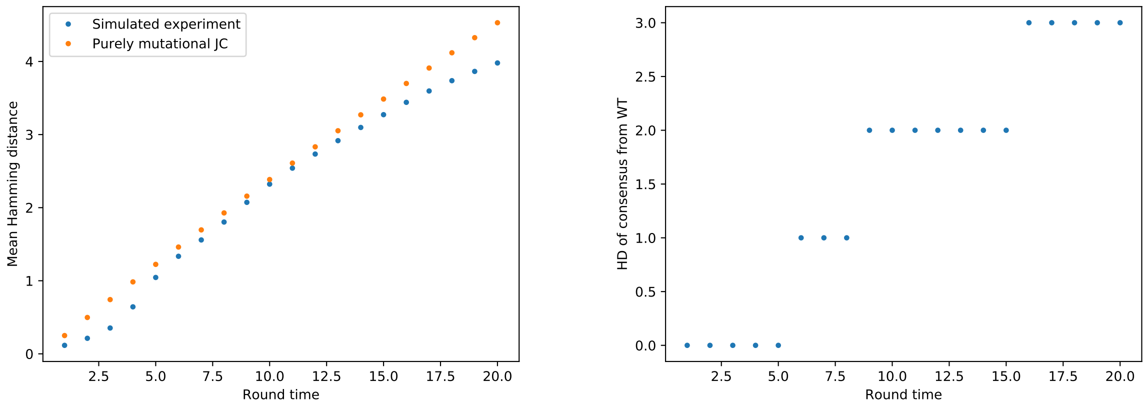


(a) (b)

Figure 3. Left: evolution of the average Hamming distance for in silico generated data (blue points), compared to the analytical prediction of the purely mutation process (orange points) with the same single site mutation probability $p = 0.01$. On the $x$ axis is the round time, whereas on the $y$ we have empirical average $\overline{\mathrm{h_D}(\boldsymbol{S}, \boldsymbol{S}^{(WT)})}$ (blue) and purely mutational theoretical average $\left\langle \mathrm{h_D}(\boldsymbol{S}, \boldsymbol{S}^{(WT)}) \right\rangle$ (orange), according to eq. (9). Right: Evolution of the HD of the consensus sequence from the wild-type. The trend displays a step behavior, remaining constant for some rounds and then increasing of a unity. Along the constant distance interval, the consensus sequence may nonetheless vary.

the pseudo-likelihood. In both cases, the inferred value is close to the ones employed to generate the synthetic data. In the panel (a) the optimal mutation probability is $p = 0.014$, to be compared with the teacher value $p = 0.01$. In general, since selection acts diminishing complexity with respect to a purely mutational process, we expect the inferred value to be smaller than the true mutation rate. However, the presence of the selection process alters the pure mutational process centered on the initial wild-type, due to the presence of distant high fitness variants the final effect on the $p$ inference is to provide a higher value than the true one.

Moreover, since $\nu(t)$ depends logarithmically on $p$, relatively small variations in the former may correspond to a relevant change of the latter. Indeed, the relative discrepancy between correct and estimated $\nu$ is much

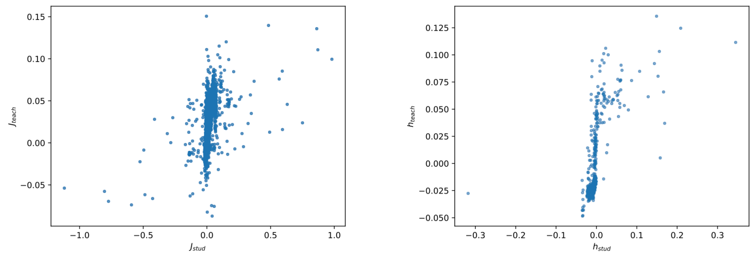

(a) $\rho_J = 0.40$, $\rho_h = 0.60$.

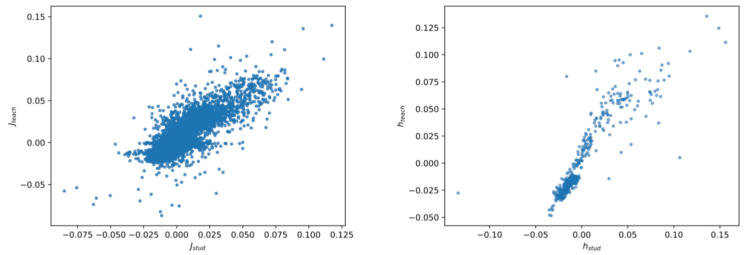

(b) $\rho_J = 0.80$, $\rho_h = 0.90$.

Figure 4. Scatter plots for selective energy parameters $\boldsymbol{J}^{(E)}$ and $\boldsymbol{h}^{(E)}$, for the same datasets reported in the main text (figure 4 panel (c)-(d)). (a) High selection low mutation regime, round 14, 16, 18, 20. (d) Low selection high mutation regime, round 2, 4, 6.

lower than between inferred $p$. This can be appreciated in table I, where we reported some values of true and inferred single site mutation probability $p$, together with their relative difference, and the normalized distance between the $\boldsymbol{\nu}$'s vectors $\|\boldsymbol{\nu}_{\text{teach}} - \boldsymbol{\nu}_{\text{stud}}\|/\|\boldsymbol{\nu}_{\text{teach}}\|$. The most off the mark inferred $p$ value coincides with the *high selection low mutation* regime. This may be ascribed to the high selective pressure characterizing this dataset.

Table I. Table reporting true and inferred value of the single site mutation probability for datasets corresponding to panels (c) and (d) of figure 4 and the first two points of figure 5 in the main text. Moreover, we reported the relative difference between the $p$'s, and the normalized distance between the vectors of Jukes-Cantor parameters $\nu$.

| Dataset | $p_{\text{teach}}$ | $p_{\text{stud}}$ | $\|\delta p\|/p_{\text{teach}}$ | $\|\|\delta\boldsymbol{\nu}\|\|/\|\|\boldsymbol{\nu}_{\text{teach}}\|\|$ |
|---|---|---|---|---|
| Fig. 5 (1,2,3) | 0.01 | 0.00435 | 0.56 | 0.12 |
| Fig. 5 (2,4,6) | 0.01 | 0.006 | 0.4 | 0.083 |
| Fig. 4 (HSLM) | 1e−3 | 1.6e−4 | 0.84 | 0.25 |
| Fig. 4 (LSHM) | 0.05 | 0.035 | 0.3 | 0.087 |

In panel (b) of figure 5, we show an example of pseudo-likelihood as a function of the $\beta$ values of two rounds. The trainig was performed on rounds 4, 8 and 12, and we infer $\beta(8)$ and $\beta(12)$ with $\beta(4) = 1$. Rescaling using $\beta(4) \equiv 1$, we expect $\beta(8) = 2$ and $\beta(12) = 3$, we obtain a maximum likelihood estimation of $(\beta(8), \beta(12)) = (2.65, 3.65)$, not far from the expected value.

In table II, we report the inferred values of the optimal inverse temperatures $\boldsymbol{\beta}$ and mutation rates $p$ for the datasets of [3] and [4], together with the corresponding regularization multipliers $\boldsymbol{\lambda} = (\lambda_J, \lambda_h) = (\lambda, 2\lambda)$.

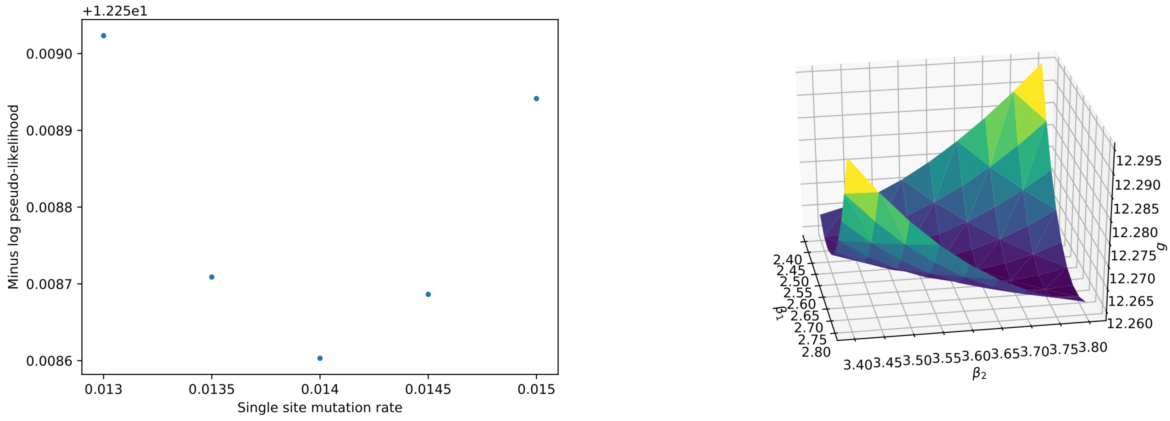


(a) (b)

Figure 5. Scan of the minus log-pseudo-likelihood as a function of the parameters $p$ (left) and $\boldsymbol{\beta}$ (right). In both cases the optimal value coincides with the minimum of the likelihood.

Table II. Values of AMaLa's time dependent parameter $\boldsymbol{\beta}$ and single site mutation rate $p$, as inferred on real directed evolution data, together with the regularization multiplier $\lambda$ (see eq. (5)).

| Protein | $\lambda$ | $\boldsymbol{\beta}_{\rm opt}$ | $p_{\rm opt}$ |
|---|---|---|---|
| PSE-1 | 0.01 | (1.0,1.7) | 0.05 |
| AAC6 | 0.005 | (1.0,1.44,1.89) | 0.05 |
| TEM-1 | 0.01 | (0.0,1.0,1.0) | 0.017 |

#### IV. SUPPLEMENTARY RESULTS ON DIRECTED EVOLUTION EXPERIMENTS

Although using the dataset described in Fantini et al. (2019)[4] to train AMaLa model provide accurate predictions of functional properties of $\beta$-lactamase TEM-1, interestingly the same is not true for contact prediction. The weak structural signal detected in the dataset was also reported by the authors of [4]. A possible reason for it is described in [5].

In figure 6 is shown the outcome of the contact prediction using the model obtain with AMaLa (over all rounds) and PlmDCA on the last round. The values of the area under the curve AUC of the specificity plot and the positive predicted value (PPV) of the first 100 predictions are: AUC(100)=0.27, PPV(100)=0.2 for DCA, and AUC(100)=0.22, PPV(100)=0.18 for AMaLa.

On the panel (a) of figure 6, it is shown the comparison between PlmDCA and AMaLa method in predicting the fitness when trained on the Fantini et al. dataset. It is reported the Pearson correlation coefficient between the model energy and the fitness score measured by Firnberg et al. (2014) [6], as processed in [7]. The score is reported as a function of the discrepancy threshold between this dataset and another independent measurement by Jacquier et al. (2013) [8] ( the same procedure it is used in [7]). More the two measures are similar less noisy they can be considered. More reliable measures have a higher correlation with the model predictions. The predictions of the AMaLa method are the same reported in figure 1 of the main text, and it is compared with the prediction of PlmDCA trained on the last round.

[1] M. Ekeberg, C. Lövkvist, Y. Lan, M. Weigt, and E. Aurell, Physical Review E **87**, 012707 (2013).
[2] M. Ekeberg, T. Hartonen, and E. Aurell, Journal of Computational Physics **276**, 341 (2014).
[3] M. A. Stiffler, F. J. Poelwijk, K. P. Brock, R. R. Stein, A. Riesselman, J. Teyra, S. S. Sidhu, D. S. Marks, N. P.

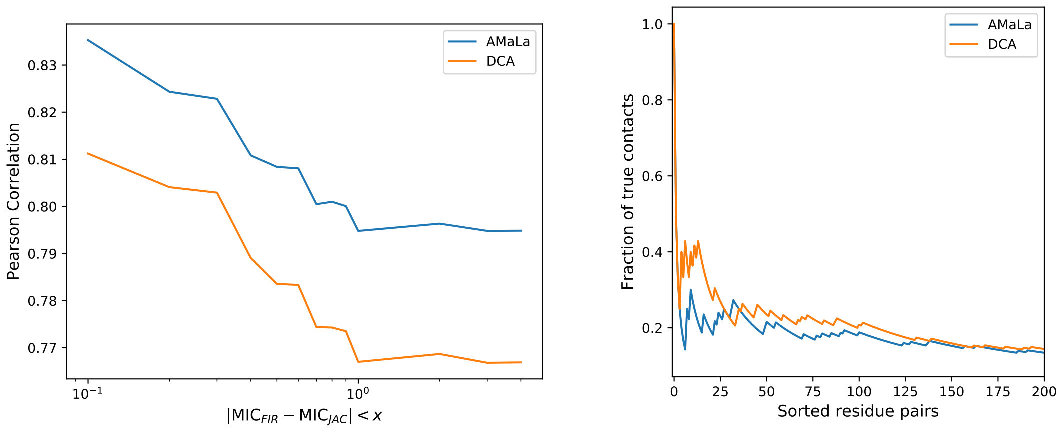


(a) (b)

Figure 6. Further results obtained on [4] dataset. Panel (a) shows a comparison between AMaLa and PlmDCA in term of fitness reconstruction, when both methods are applied to the data of [4]. On panel (b) we reported the sensitivity plot for contact prediction obtained from both inference methods, which contrarily to fitness reconstruction, seems to bring about a low information content.

Gauthier, and C. Sander, Cell Systems **10**, 15 (2020).
